# Bovine neutrophils release extracellular traps and cooperate with macrophages in *Mycobacterium avium* subsp. *paratuberculosis* clearance *in vitro*

**DOI:** 10.1101/2020.12.30.424791

**Authors:** I Ladero-Auñon, E Molina, A Holder, J Kolakowski, H Harris, A Urkitza, J Anguita, D Werling, N Elguezabal

## Abstract

*Mycobacterium avium* subsp. *paratuberculosis* (Map) is the underlying pathogen causing bovine paratuberculosis (PTB), an enteric granulomatous disease that mainly affects ruminants and for which an effective treatment is needed. Macrophages are the primary target cells for Map, which survives and replicates intracellularly by inhibiting phagosome maturation. Neutrophils are present at disease sites during the early stages of the infection, but seem to be absent in the late stage, in contrast to healthy tissue. Although neutrophil activity has been reported to be impaired following Map infection, their role in PTB pathogenesis has not been fully defined. Neutrophils are capable of releasing extracellular traps consisting of extruded DNA and proteins that immobilize and kill microorganisms, but this mechanism has not been evaluated against Map. Our main objective was to study the interaction of neutrophils with macrophages during an *in vitro* mycobacterial infection. For this purpose, neutrophils and macrophages from the same animal were cultured alone or together in the presence of Map or *Mycobacterium bovis* Bacillus-Calmette-Guérin (BCG). Extracellular trap release, mycobacteria killing as well as IL-1β and IL-8 release were assessed. Neutrophils released extracellular traps against mycobacteria when cultured alone and in the presence of macrophages without direct cell contact, but resulted inhibited in direct contact. Macrophages were extremely efficient at killing BCG, but ineffective at killing Map. In contrast, neutrophils showed similar killing rates for both mycobacteria. Co-cultures infected with Map showed the expected killing effect of combining both cell types, whereas co-cultures infected with BCG showed a potentiated killing effect beyond the expected one, indicating a potential synergistic cooperation. In both cases, IL-1β and IL-8 levels were lower in co-cultures, suggestive of a reduced inflammatory reaction. These data indicate that cooperation of both cell types can be beneficial in terms of decreasing the inflammatory reaction while the effective elimination of Map can be compromised. These results suggest that neutrophils are effective at Map killing and can exert protective mechanisms against Map that seem to fail during PTB disease after the arrival of macrophages at the infection site.

## 1 Introduction

*Mycobacterium avium* subsp. *paratuberculosis* (Map) is the aetiological agent of paratuberculosis (PTB; Johne’s disease), a chronic granulomatous enteritis of ruminants characterized by cachexia and severe diarrhoea. It results in production losses due to sub-clinically infection and animal losses due to clinically active disease (1). Infected animals in the subclinical stage shed Map intermittently to the environment, making its control difficult. Vaccination against PTB has been shown to reduce shedding and therefore in-herd transmission (2) (3). However, vaccination does not provide complete protection from infection (4) and due to the nature of the Map antigens interferes with the official tests for bovine tuberculosis (bTB) eradication program.

Classically, PTB research has been focused on the action of macrophages (MΦ) as the main target cells. After infection, Map is able to survive and replicate intracellularly by inhibiting phagosome maturation. However, less attention has been paid to the role that polymorphonuclear neutrophils (PMN) may play in this process, although their presence is undeniable especially during the early stages of the infection (5). PMNs are short-lived effector cells capable of phagocytosing pathogens, killing microbes through the production of reactive oxygen species (ROS), lytic enzymes with potent antimicrobial activity (6) as well as the release of extracellular traps (ETs) (7, 8).

ET formation has been demonstrated as an important novel effector mechanism against pathogens in PMNs (8) and MΦs (9). ETs are extracellular structures composed of chromatin and granule proteins that trap and kill microorganisms (10). Their formation is triggered by pathogens, danger associated molecular patterns, exogenous compounds, platelets, antibodies and inflammatory cytokines such as IL-8 and TNF (11, 12). MΦ have been reported to produce ETs in response to *Staphylococcus aureus* (13), *Mycobacterium tuberculosis* (Mtb) (14) and the parasite *Besnoitia besnoiti* (15). However, this antimicrobial mechanism does not seem to be dominant in MΦ. It is more likely that the main function of MΦ in this context is to clear ETs produced by PMNs and the resulting apoptotic PMNs rather than to produce ETs themselves (16).

Indeed, several studies have described the cooperation of PMN and MΦ to remove pathogens at the inflammation site (17, 18), with apoptotic PMNs being phagocytosed by MΦ’ in a process called efferocytosis, which can potentially result in cross-presentation. This mechanism provides MΦ with antimicrobial molecules that are lacking in mature MΦ and that aid the killing of intracellular pathogens (18, 19). PMN also transfer ingested intracellular pathogens to MΦ during efferocytosis (18).

In addition to PMN and MΦ, the pro-inflammatory cytokines IL-1β and IL-8 are also essential components in the formation of granulomas in mycobacterial infections. IL-1β is a key mediator of the innate inflammatory response that attracts further MΦ (20) and leads to a general activation of the immune system. Furthermore, IL-1β seems to play an important role in protecting mice against experimental Map infection (21). Although essential for resistance to infection, IL-1β also exacerbates damage during chronic diseases and acute injuries. Its induction by Map may result in attraction of MΦ to the site of infection, thus ensuring its survival and dissemination (22), and this process may further be aided by the release of IL-8 release by PMN. IL-8 plays an important role in leukocyte recruitment, granuloma formation, and respiratory burst in response to *Mycobacterium tuberculosis* (Mtb), leading to enhanced phagocytosis and killing of Mtb (23), and it is considered as a major ET inducer for PMN (24).

There is little research on the protective role of PMN in Map infection in cattle. PMN isolated from cattle infected sub-clinically with Map showed diminished migratory properties when stimulated *in vitro* compared to PMN of healthy cattle (25), suggesting that Map infection potentially undermines PMN functionality. Indeed, PMN seem to be absent in tissues from cows with clinical PTB (26). In the last years, several transcriptome studies have supported the idea that Map infection causes an impairment of PMN recruitment (27, 28) and the downregulation of antimicrobial peptide production by PMN, such as those of beta-defensins (29) and cathelicidins (28). A recent study has shown that cathelicidin LL-37 produced by human PMN restricts Mtb growth (30) and facilitates Map clearance in murine MΦ by suppressing the production of tissue-damaging inflammatory cytokines such as IL-8, TNF and IFN-γ (31). Taken together, these studies suggest that PMN may play a more important role than previously thought during the early stages of Map infection. Thus, studying this immune cell type during Map infection may help to identify novel intervention strategies.

In order to gain knowledge on PMN and Map, we studied the interaction of bovine monocyte-derived MΦ (MDM) and PMNs, either alone or in co-cultures, infected with Map and BCG, by evaluating ET formation, bacterial killing and pro-inflammatory cytokine production. To our knowledge, this is the first study to investigate ET release in response to Map exposure.

## 2 Materials and Methods

### 2.1 Animals

Blood samples were drawn from healthy Holstein Friesian cows of a commercial dairy farm located in the Basque Country. The herd is enrolled in the national bTB eradication program. Animals on the farm tested negative for bTb by skin-test over the last five years. All animals used in this study (n=4) tested negative for Map shedding in faeces, as confirmed by PCR, and were negative for Map-specific antibodies to Map, as determined by PPA-3 ELISA.

Animals used in this study were submitted only to procedures that according to European (Directive 2010/63/EU of the European Parliament and of the Council of 22 September 2010 on the protection of animals used for scientific purposes. Chapter 1, Article 1, Section 5, paragraphs b and f) and Spanish (Real Decreto 53/2013, de 1 de febrero, por el que se establecen las normas básicas aplicables para la protección de los animales utilizados en experimentación y otros fines científicos, incluyendo la docencia, Article 2, Section 5, Paragraphs b and f) legislation on experimental animals are exempt from its application. The animals, belonging to a registered commercial farm supervised by the local livestock authority (Servicio de Ganadería de la Diputación Foral de Bizkaia) were submitted only to the introduction of a needle in accordance with good veterinary practice and were not killed in relationship with this study.

### 2.2 Bovine peripheral blood monocyte isolation and generation of macrophages

Blood was taken from the jugular vein using a 16G x 11/2 hypodermic needle into a blood-collection bag containing 63 mL of citrate phosphate dextrose adenine (CPDA; TerumoBCT Teruflex) for a total capacity of 450 mL, which was used for monocyte isolation. One month later, 40 mL of blood from the same cows were collected with a 18G hypodermic needle into a 50 mL tube containing 7 mL of CPDA for PMN isolation. Blood samples were processed within two hours of the extraction.

Peripheral blood mononuclear cells (PBMCs) were separated using Histopaque 1077® and monocytes were selected using mouse anti-human CD14-coupled microbeads and MS columns (Miltenyi) following the manufacturer’s recommendations. Greater than 90% of purity of monocytes was achieved as determined by flow cytometry using a mouse anti-bovine CD14 FITC-labelled antibody (BioRad).

For the production of monocyte-derived macrophages (MDM), monocytes were plated at 1×10^5^ cells/well for co-cultures and at 2×10^5^ cells/well for MDM cultures in wells of 96-well plates for fluorimetric assays to quantify ET release. For immunofluorescence microscopy, monocytes were plated at 1×10^5^ cells/well for co-cultures and at 2×10^5^ cells/well for MDM cultures on 16-well chamber slides (178599PK Thermo Scientific). For contact independent cultures, monocytes were plated at 5×10^5^ cells/well for co-cultures and at 1×10^6^ cells/well in wells of 24-well plates. During maturation to MΦ, monocytes were cultured in RPMI 1640 media without phenol red (Gibco 11835030), supplemented with 2mM L-glutamine, 10% foetal bovine serum (FBS), 1% penicillin-streptomycin and 40 ng/mL recombinant bovine M-CSF (Kingfisher Biotech, Inc. RP1353B) at 37°C and 5% CO2. The media was replaced every three days and prior to the beginning of the mycobacterial infection assays. After 7 days of culture, MΦ morphology was confirmed by light microscopy (increased size, increased adherence, cytoplasmic granularity and presence of pseudopods).

### 2.3 Bovine Neutrophil isolation

A week after PBMC isolation, 25 mL of blood were drawn from the same cows by venepuncture and mixed with 25 mL of PBS supplemented with 2% FBS. 25 mL of the mixture were layered on top of 15 mL of Histopaque® 1077 (Sigma) and centrifuged at 1200 x g for 45 min at RT with no brakes. PMNs were located at the lower end of the tube and were subjected to two flash hypotonic lysis using 25 mL of sterile distilled water during 20-30 sec, followed by 2.5 mL of a 10% NaCl solution and topped up with PBS-2% FBS, in order to eliminate remaining red blood cells. PMNs were then resuspended in RPMI 1640 with glutamine and without phenol red and counted. Over 90% purity was achieved as determined by FSC and SSC gating and visualization of characteristic nuclei after DAPI staining under a fluorescence microscope.

### 2.4 Bacterial strains and culture

Map strains K10, Map K10-GFP (kindly provided by Dr. Jeroen de Buck, University of Calgary, (32)), *M*.*bovis* BCG Danish strain 1331 and *M. bovis* BCG Danish strain 1331-GFP (32) were used in the assays. All strains were grown to exponential phase at 37°C under aerobic conditions. Map-GFP and BCG-GFP were cultured for three weeks on Middlebrook 7H9 OADC with Kanamycin (25μg/mL) to select for GFP-plasmid carriers and supplementation with Mycobactin J was added for Map. Map strain K10 and BCG Danish strain 1331 were used for ET isolation and quantification and were cultured in the same medium without kanamycin. Cultures were adjusted after reaching an OD of 0.7 (3×10^8^ bacteria/mL) and colony forming units (CFU) were confirmed on agar after using 10-fold serial dilutions of bacterial suspensions. In experiments comparing the response to live and killed bacteria, heat-inactivation was performed by heating bacteria for 30 min at 85°C. Inactivation of bacteria was confirmed by plating on agar plates.

### 2.5 Cell cultures and *in vitro* infection

PMNs and MDMs isolated from the same cow were cultured separately or co-cultured together in the same well (CC; see Figures 1A and 3A for details). In order to study cellular changes following paracrine signalling in the absence of cell-to-cell contact between PMNs and MDMs as well as to determine ET formation, a transwell co-culture system was also employed. PMNs were seeded in the transwell insert (pore diameter 0.4 μm; Thermo Fisher Scientific 140620) to generate insert PMNs (TW-PMNs), whereas MDMs were seeded into the lower well (TW-MDM). To assess the need for direct cell contact, conditioned MDM media (TW-CM) was produced by incubating MDMs with Map, BCG and Zymosan for 4h at 37°C and 5% CO_2_, which was subsequently added to the lower well underneath the insert containing PMNs (TW-PMN-CM). ET release quantification and mycobacteria killing were performed on 24 and 96 well plates. ET visualization was performed on 16-well chamber slides (Nunc Labtek).

**Figure 1.**
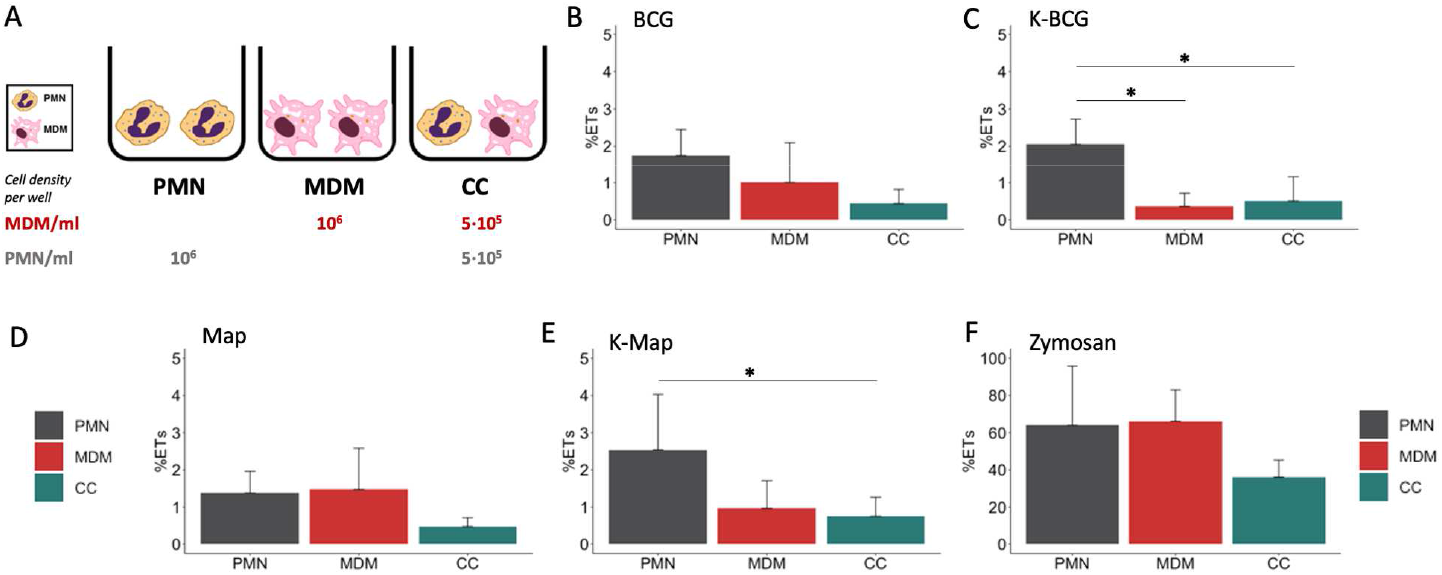
Fluorimetric ET release quantification of PMNs, MDMs and CCs. (A) Cell culture types and conditions in 96 well plates. ET release quantified by fluorimetry of cultures stimulated with (B) BCG, (C) K-BCG: killed BCG, (D) Map, (E) K-Map: killed Map and (F) Zymosan. Data are composed of combined values obtained of all cows (n=4), with samples run in triplicates. * depicts a p value of p < 0.05.

Once cell cultures were set, these were stimulated with live or heat-inactivated Map K10-GFP, *M. bovis* BCG -GFP, Map K10, *M. bovis* BCG at either a MOI of 1 (ET release) or a MOI of 5 (survival assay), zymosan (1mg/mL; Sigma Z4250), or left untreated (negative control). The original bacterial strains with no GFP were used for the isolation and quantification of NETs in the fluorometric assay to avoid GFP fluorescence interference.

Cultures were incubated at 37°C and 5% CO_2_ for 4h for ET release evaluation and for 24h for bacteria killing and cytokine levels.

### 2.6 Isolation and quantification of ETs

ET release was assayed as described by Köckritz-Blickwede et al (33) with some modifications. Briefly, ETs generated by cells were digested with 500 mU/mL of micrococcal nuclease (10107921001 Roche) for 10 mins at 37°C 5% CO2. The nuclease activity was stopped by the addition of 5 mM EDTA; thereafter, culture supernatants were collected and stored at 4°C overnight. Total DNA was extracted from naive cells with DNeasy® blood & tissue kit (Qiagen) following the manufacturer’s instructions. Extracted DNA was solubilized in TE buffer. Both ETs and genomic DNA was quantified using Quant-iT™ PicoGreen™ assay (Thermofisher) according to the manufacturer’s instructions. Plates were read in a fluorescence microplate reader (Synery HTX, Biotek) with filter settings at 488 nm excitation and 520 nm emission. The percentage of DNA released as ET-DNA was calculated by dividing the amount of isolated ET-DNA by the total amount of genomic DNA.

### 2.7 Visualisation and quantification of ET by immunofluorescence microscopy

To assess histone staining as a marker for ET formation, cells were seeded into 16-well chamber slides for subsequent antibody staining and microscopy according to Conejeros et al (34). After 4 h of incubation with either Map-GFP, BCG-GFP or zymosan, cells were fixed for 15 min with 4 % formaldehyde. Cells were washed twice with PBS and kept in PBS at 4°C until staining was performed. Subsequently, cells were permeabilized for 15 min with PBS 0.1% Triton X-100, blocked with 1% goat serum containing 0.05% Tween 20 and 3% BSA-PBS. The immunofluorescence staining was performed overnight at 4°C using a mouse anti-pan-histone antibody (Merk MAB3422; diluted 1:200). Thereafter, cells were washed three-times with PBS and incubated with anti-mouse Alexa Fluor 594 labelled antibody (Invitrogen A-1105, diluted 1:500) for 30 min at RT. Following further three washes with PBS, wells were detached from the slides and mounting medium containing DAPI was added before coverslips were added. Preparations were let dry and observed at 400x magnification on a DMi8 fluorescence inverted microscope (LEICA®). Pictures were taken using a DFC3000 G camera coupled to the microscope.

Quantification was performed by taking snapshots of five fields containing visible ETs with histone staining. Analysis was performed using the Image J software package. Snapshots in 8-bit format were analysed using the following pipeline. Threshold was adjusted depending on the micrograph, being 4 for low level and 255 for high level. The percentage of picture area occupied by ETs plus nuclei (total DNA) was measured by adjusting the settings for fluorescent particle size: 0-infinity and circularity: 0-1. The percentage of picture area occupied by nuclei were analysed adjusting the settings for particle size: 200-5000 and circularity: 0.3-1. ET percentage was calculated as total DNA percentage minus nuclei percentage. In the case of cultures that did not contain ETs, nuclei were measured, and ET release was considered zero.

### 2.8 Mycobacteria killing assay

PMNs, MDMs and co-cultures inoculated with BCG -GFP and Map-GFP were prepared. At time 0 (4°C control) and 24 h, the supernatants were removed and centrifuged to pellet non-internalized mycobacteria. Supernatants were stored for cytokine analysis. Wells were washed twice with PBS to remove the remaining non-internalized bacteria. Bacterial pellets and washes of remaining bacteria were centrifuged together, and the final pellets were resuspended in 100 μL of filtered PBS. Adherent cells were lysed by vigorous pipetting with 0.5 mL of 0.1% Triton X-100 (Sigma-Aldrich) in sterile water for 10 min at RT.

Two serial 10-fold dilutions of each lysed sample and supernatant were performed in a total volume of 1 mL using filtered PBS. 200 μl of each dilution was inoculated in duplicate in Middlebrook 7H9 OADC-Kanamycin (25μg/mL) agar plates, supplemented with mycobactin J in the case of Map cultures. Seeded agar plates were allowed to dry at RT until humidity was no longer visible. Plates were sealed with tape to avoid desiccation and incubated at 37 ± 1 °C for 6 weeks. CFU were counted and 0h (4°C control) CFU were considered total inoculated bacteria, 24h CFU supernatant bacteria were considered non-internalized and non-killed and 24h CFU cell bacteria were considered internalized and non-killed bacteria. Killed bacteria were estimated as total inoculated bacteria minus (24h CFU supernatant bacteria plus 24h CFU cell bacteria). CFU counts were multiplied by the inverse dilution factor and by the seeded volume. The mean of duplicate plates was calculated and the group mean for each culture type was calculated. Supernatants’ fraction and cell-lysis’ fractions were divided by the 100 % survival control to calculate the survival percentage of each portion and the killing rate.

### 2.9 Cytokine detection

Commercially available direct ELISA kits were performed to determine bovine IL-1β (ESS0027 Invitrogen®) and IL-8 (3114-1A-6 MABTECH) release in 24h culture supernatants using the provided manufacturer’s instructions. IL-1β ELISA is based on streptavidin-HRP and IL-8 on streptavidin-ALP. Absorbances were measured using an automated ELISA plate reader (Multiskan EX®, Thermo Lab Systems, Finland) and in all cases standard curves were used to determine each cytokine amount in the supernatant samples.

### 2.10 Statistics

Data were assessed for normal distribution and homoscedasticity using Shapiro-Wilk and Bartlett’s test, respectively. All data are presented as the mean +/-standard deviation (SD), with samples being analysed in triplicates (ET release by fluorimetry) or duplicates (bacterial killing and cytokine release). For ET release, percentage differences between values obtained for the mean of each culture type were calculated using pairwise comparisons (t-tests) with pooled SD for all the stimuli except for values obtained for unstimulated and zymosan stimulated cultures. Here, values were not normally distributed and thus comparisons were calculated using non-pooled SD test.

The number of bacteria (expressed as CFU) in the inoculum (0 h) was considered as 100% viable bacteria. The number of bacteria being alive after 24 h incubation was calculated as the sum of internalized mycobacteria and mycobacteria being present in the supernatant, with the killing rate being the difference between the bacteria at 0h and the survival at 24 h, expressed as percentage. Mean values obtained for killing rates under different conditions were compared by multiple pairwise comparisons using t tests with non-pooled SD with Benjamini-Hochberg correction. To obtain killing rates for co-cultures, values obtained for the killing rate of individual cultures (PMN, MDM) were divided by two and obtained values were added to each other.

Differences in IL-1β levels were compared by Kruskal-Wallis test with the post-hoc Dunn’s multiple comparison test. IL-8 level differences were calculated through multiple pairwise comparisons using t tests with pooled SD.

To identify correlations between variables, Pearson (ρ_x,y_) and Spearman (ρ) coefficients were calculated to detect linear and non-linear correlations and in each case the best fitting coefficient value was chosen.

All statistical tests were performed using R studio desktop (version 1.2.5033; (RStudio Team, 2019. RStudio: Integrated Development for R. RStudio, Inc., Boston, MA URL http://www.rstudio.com/), and a p-value of < 0.05 was considered statistically significant.

## 3 Results

### 3.1 PMNs release extracellular traps against both mycobacteria killed and alive in contrast to MDMs

In a first set of experiments, we assessed the induction of ET in PMNs and MDMs, either alone or in co-cultures, in response to Map and BCG. In order to assess whether this mechanism would rely on live or heat-killed bacteria, all culture types were incubated with both live and heat-inactivated Map and BCG. Fluorometric analysis revealed that both cell types cultured individually extruded similar detectable DNA levels in response to zymosan (Figure 1F), whereas lower ET levels were detected when cells were cultured together (Figure 1F). Levels of ET formation in response to either live mycobacteria were similar for all culture cell type (Figure 1B and D), and lower compared to cells incubated with heat-killed mycobacteria. For these conditions, PMNs showed higher ET release against killed mycobacteria (Figures 1C and E) compared to MDMs (K-BCG, p=0.03; K-Map, p=0.065) and to CC (K-BCG, p=0.04; K-Map, p=0.043).

### 3.2 PMNs and MDMs alone or in direct co-cultures exert different effector mechanisms against BCG and Map

Although fluorometric quantification analysis revealed significant differences in ET release among cell culture types, further analyses were performed to discriminate between DNA from ET release and DNA liberated by other means. Image analysis quantification of ETs showed that PMNs liberated significantly higher levels of ETs (6.52% - 7.23%) compared to MDMs (0%) and CCs (1.13% - 2.03%) and MDMs (0%) when stimulated with both mycobacteria, independent of whether the bacteria were alive or heat-killed (Figure 2A-D). As seen before, MDMs produced ET-like structures only when stimulated with zymosan (Figure 2E). MDMs and PMNs, cultured either alone or in co-cultures and stimulated with zymosan lost normal nuclei structure, ending in nuclear DNA extrusion (Figure 2E). However, co-cultures seemed to contain fewer ETs, which could be due to MDMs phagocytosing liberated ETs by PMNs or because direct contact of both cell types inhibits ET release.

**Figure 2.**
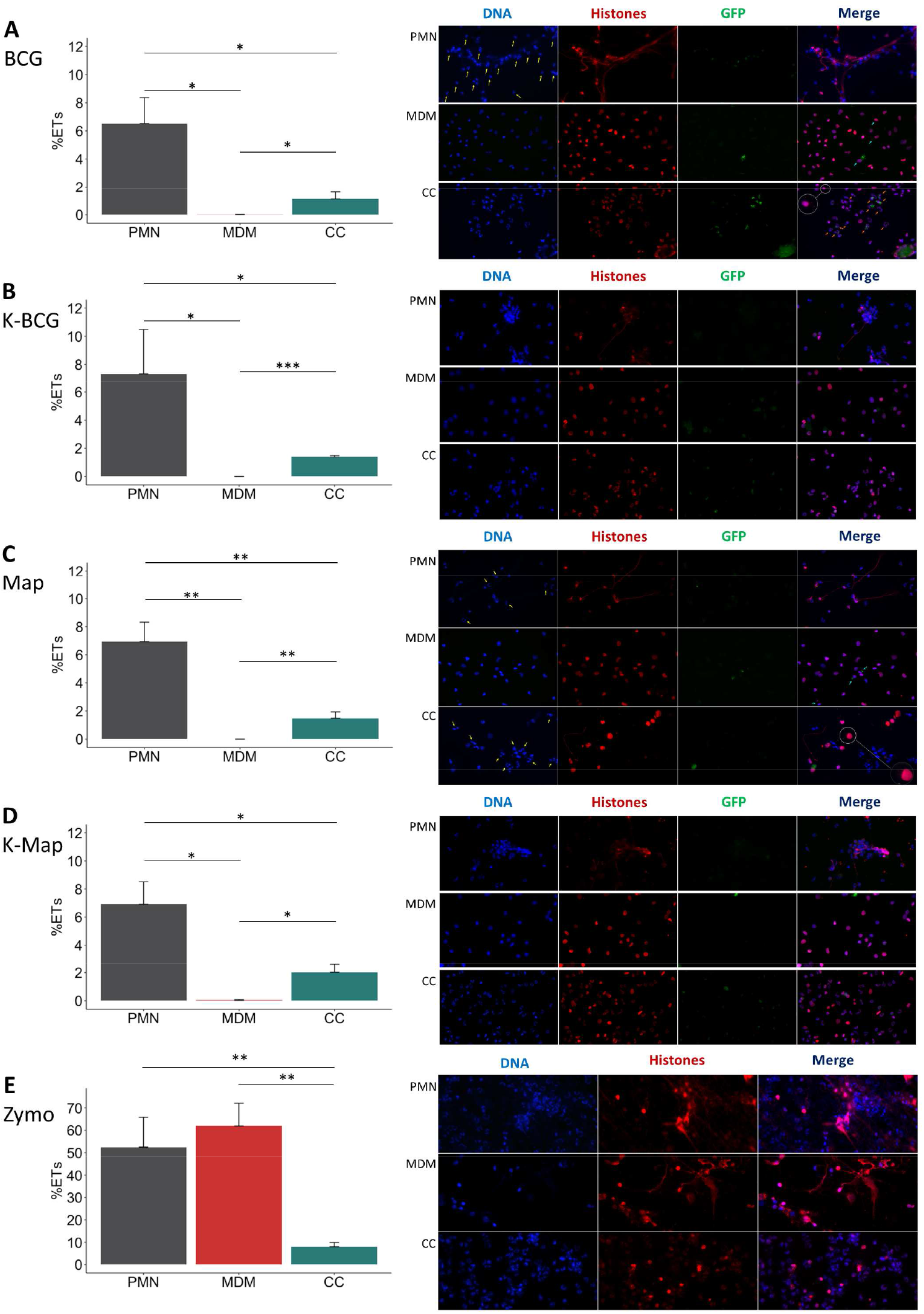
ET release analysis of microscope images. In the left of each section bar plots representing ET release quantified by immunofluorescence microscopy image analysis and in the right representative corresponding examples of micrographs (40x) of PMNs, MDMs and CCs stimulated with (A) BCG, (B) K-BCG: killed BCG, (C) Map, (D) K-Map: killed Map and (E) Zymosan. DAPI stained DNA in blue, histones in red and BCG-GFP or Map-GFP in green. Yellow arrows indicate nuclei stained with DAPI, which are absent in the red channel corresponding with anti-histone immunolabeling. Light-blue arrows indicate MDM phagocytosis. Orange arrows indicate PMN phagocytosis. A white circle has an augmented image showing a red punctate pattern in MDM cytoplasm. Data composed of combined values obtained of all cows (n=4), with samples run in duplicate and 5 fields quantified per micrograph. (* p<0.05; ** p<0.01;*** p<0.001)

The microscopic evaluations also revealed that ETs released by PMN were the main source of histone-staining (shown in red), as this staining was mainly within the extruded ETs and not always associated with DAPI-stained nuclei (Figure 2A and C; PMNs and PMNs and CC). In agreement with data shown in Figure 1, MDMs showed internalized mycobacteria but did not show ET release (Figure 2A and C, MDMs), and anti-histone staining was always associated with the nuclei (as recognised by DAPI staining), independent of bacterial preparation used to stimulate MDM (Figure 2A-D; MDM). In addition, it seemed that MDMs showed less mycobacterial internalization when co-cultured with PMN, whereas the opposite was observed with PMNs (Figure 2A and C; CC). However, in Map stimulated CC, PMNs nuclei were not always labelled by the anti-histone antibody, and many PMNs showed condensed nuclei, potentially indicative of apoptosis (Figure 2C; CC). Furthermore, ETs were rarely detected and MDMs presented a red punctate pattern in the cytoplasm (Figure 2A and C; CCs) which could support the hypothesis that MDM phagocytosed histones liberated by PMNs.

### 3.3 ET release by PMNs in response to mycobacteria is not affected by stimulated MDMs soluble factors

Having established that both mycobacterial species induce ET formation in PMNs, but less so in MDMs or co-culture of both cells, we next assessed if ET release was influenced by secreted mediators or required direct cell-cell contact (Figure 3A). To further address the possible paracrine effect of MDMs, PMNs were also cultured in the transwell system (TW-PMN-CM) containing macrophage-conditioned media (CM) (Figure 3B). Additional controls included unstimulated and stimulated PMN kept in the transwell insert on their own (TW-CTRL and TW-PMN, respectively) to exclude an impact on ET release by the transwell culture condition compared to PMNs kept in the lower well (PMN) (see Figure 3A).

**Figure 3.**
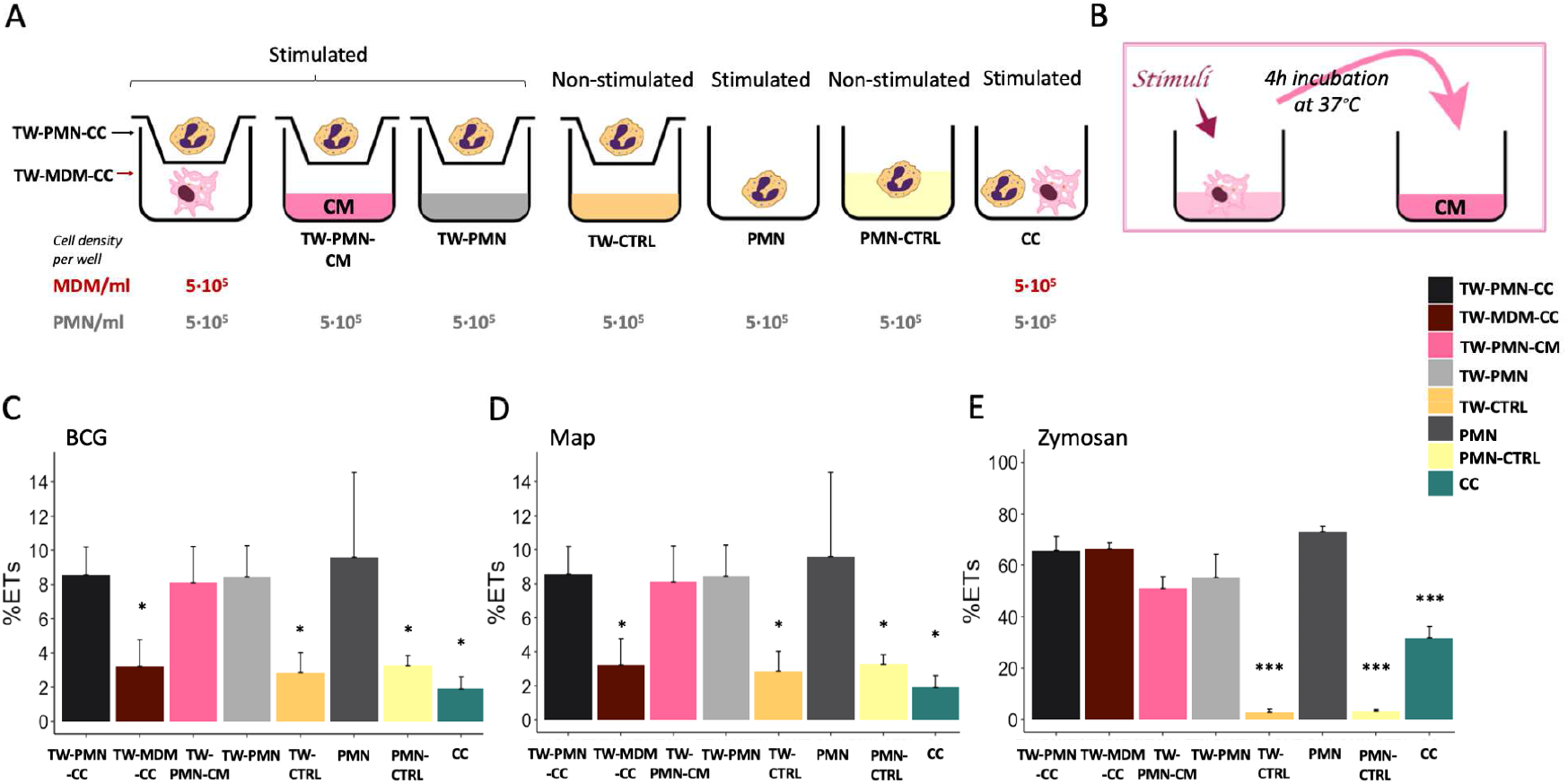
ET release quantified by fluorimetry of the transwell system experiment. (A) Culture types and conditions in 24 well plates. (B) Conditioned media preparation from stimulated macrophages for PMN stimulation. Fluorimetric quantification of ET release of (C) BCG, (D) Map and (E) Zymosan stimulated cultures. Data composed of combined values obtained of all cows (n=3), with samples run in duplicate (* p<0.05, **; p<0.01; *** p<0.001). TW: transwell, CC: co-culture, CM: conditioned media, CTRL: non-stimulated control.

PMN incubated either in the transwell insert (TW-CTRL) or directly in the well (PMN-CTRL) showed no differences in baseline ET release, indicating that the culture conditions did not impact on ET release.

PMNs released equivalent levels of ETs in response to both mycobacteria when cultured either alone in the lower well (PMN) in the presence of MDM conditioned media (TW-PMN-CM), or cultured in the insert of the transwell (TW-PMN) (Figure 3C and D). In contrast, MDMs do not release ETs in transwell co-culture with PMNs when stimulated with mycobacteria (Figure 3C and D), confirming results obtained previously.

Direct cell contact induced the lowest ET release compared to the rest of the co-culture types independent whether cells were stimulated with either mycobacteria (p<0.05), and this effect was also present when cells were stimulated with zymosan (Fig. 3E). Overall, the response to zymosan followed the response seen to either mycobacteria with the exemption of MDM that sowed a similar ET release compared to PMN (60-70%) upon stimulation with zymosan.

### 3.4 PMNs show similar killing levels of Map and BCG, whilst MDMs are more effective at killing BCG

After demonstrating that ET release can be triggered by mycobacteria after 4 hours, we next assessed whether ET formation at an early time point may be beneficial in terms of bacterial clearance. For this purpose, the killing capacity of PMNs, MDMs and co-cultures against both mycobacteria after 24 h was quantified. Total inoculated CFU (0 h) and CFU recovered at 24h for all cultures are shown in Figure 4A and C. MDMs alone and CCs were more effective at killing BCG compared to PMNs alone, whereas PMNs were more effective at killing Map compared to MDMs.

**Figure 4.**
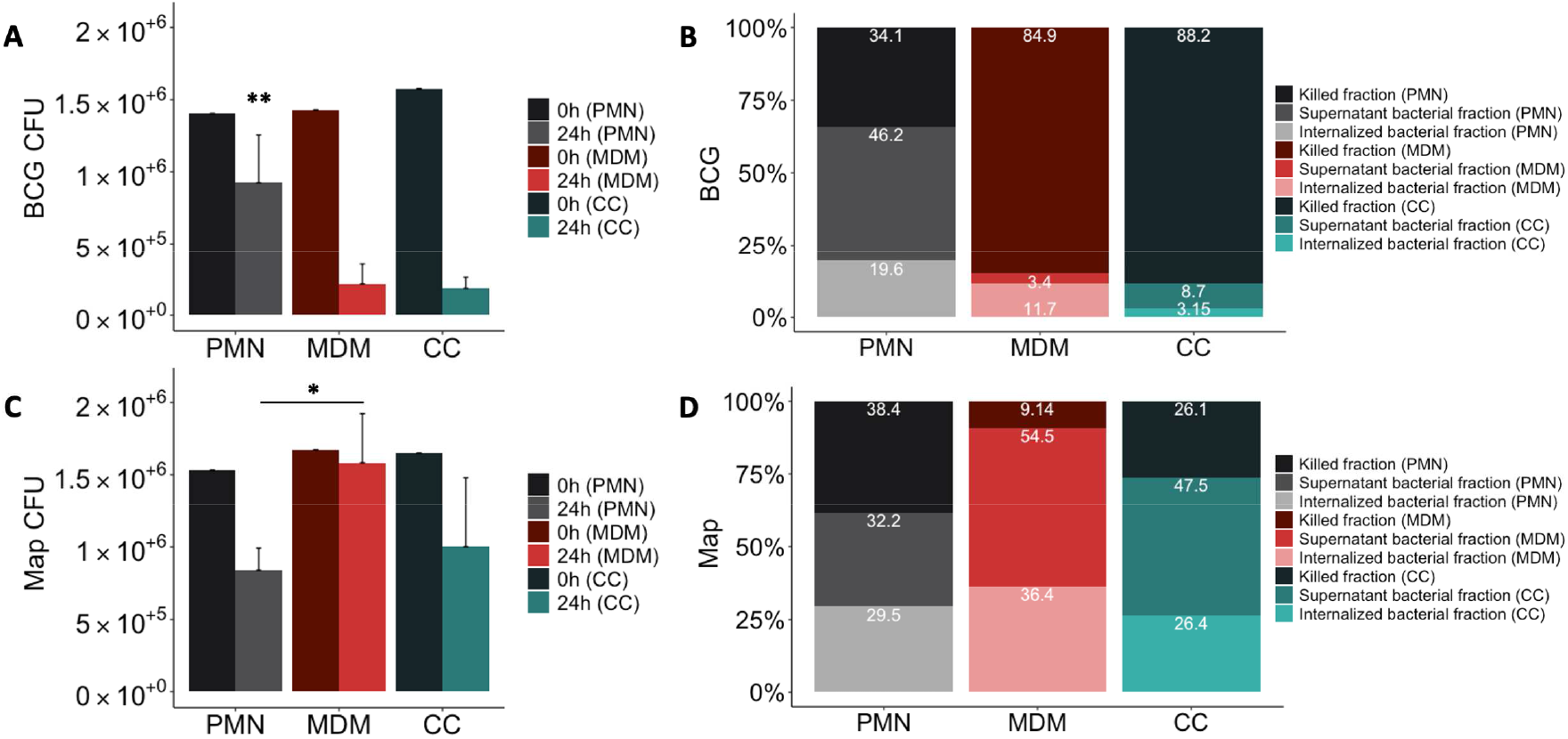
Mycobacterial killing assay results. Distribution of mycobacteria (A) total CFUs of BCG-GFP, (B) % BCG-GFP, (C) total CFUs Map-GFP and (D) % Map-GFP in 24h cultures of PMNs in grey, MDMs in red and in direct contact co-cultures (CC) in green. In (A) and (C) in the dark colour the 0h control and in the light colour the 24h assay of the corresponding culture type. In (B) and (D) in the darkest colours the fraction of killed bacteria, in the intermediate intensity colours the fraction of live bacteria non-internalized from the supernatant and in the lightest colours the proportion of internalized bacteria that survived inside the adherent cells of the corresponding culture type. Data composed of combined values obtained of all cows (n=4), with samples run in duplicate (* p=<0.05, **, p=<0.01).

The resulting distribution of live and killed BCG and Map in culture cells and supernatant is shown in Figure 4B and D. Statistical analysis revealed significant differences between MDMs, PMNs and CCs showing higher killing of BCG in MDM cultures compared to PMNs (84.85% vs 34.15%, p=0.0016), and higher Map killing in PMN cultures than in MDMs (38.36% vs 9.14%, p=0.045). PMN culture killing rate against BCG was lower than in co-cultures (34.14% vs 88.19%; p=0.001), whereas MDM culture killing rate against Map was lower compared to co-cultures (9.14% vs 26.10%; p=0.065), although it was not statistically significant. Considering that CCs include half of the amount of each cell type, and that each cell type shows a certain killing capacity in individual culture, a theorical killing rate for CCs against each mycobacterium was calculated by dividing each individual culture (PMN, MDM) killing rate by two and adding both obtained values. Results indicate that co-cultures infected with BCG showed improved killing results (expected killing: 59.49% vs observed killing: 88.19%) than individual cultures. In contrast, this synergic effect was not evident in co-cultures infected with Map, showing only 2.52% of improvement (expected killing: 23.58 % vs observed killing: 26.1%).

When bacterial survival inside adherent cells of each cell-culture type was compared (Figure 4B and 4D), significant differences were not observed for either BCG or Map. However, non-internalized BCG survival in PMN cultures (46.23%) was higher than in MDM cultures (3.44%, p=0.021) and in co-cultures (8.65%, p=0.021), whereas non-internalized Map survival in PMN cultures was lower than in MDM cultures (54.46% vs 32.17%, p=0.03). Map-GFP survival inside adherent cells was higher than BCG-GFP as expected.

### 3.5 MDMs alone release higher levels of IL-1β and IL-8 compared to PMNs and alone or co-cultured with PMNs in response to both mycobacteria

Having assessed the killing capacity of both cell types to different mycobacterial strains, we next investigated whether their cytokine response follows a similar pattern.

Considering that IL-8 is a major ET inducer in PMNs (23) and IL-1β is a key mediator of the inflammatory response that attracts MΦ to the infection site (19), we analysed the levels of both cytokines in the supernatant. IL-1β release after 24h showed significant differences between the cells involved as well as in response to both mycobacteria.

MDM cultures released higher concentrations of IL-1β (Figure 5A) in response to BCG compared to PMN (p<0.001) and CC (p=0.009) and in response to Map compared to PMN (p=0.035). Furthermore, MDM released significantly more IL-1 β in response to BCG compared to Map.

**Figure 5.**
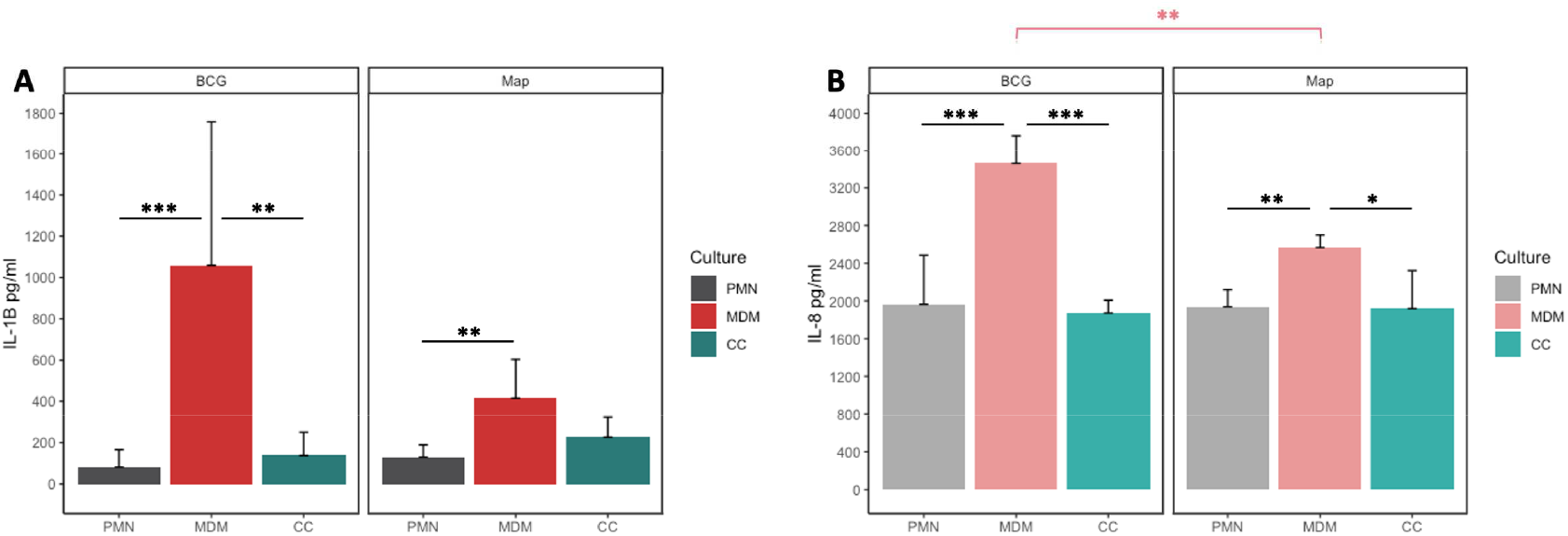
(A) IL-1β and (B) IL-8 release of PMN, MDM and CC stimulated with BCG and Map after 24h. Data composed of combined values obtained of all cows (n=4), with samples run in duplicate (* p<0.05, **; p<0.01; *** p<0.001).

Similar as seen for IL-1β, MDM secreted more IL-8 into the supernatant in response to both mycobacteria (Figure 5B) compared to co-cultures (BCG; p< 0.001, Map; p=0.013) or PMNs alone (BCG; p=0.001, Map; p= 0.005). As before, MDMs challenged with BCG produced significantly more IL-8 than challenge with Map (p=0.011).

### 3.6 Correlation analysis of cytokine production and bacteria killing rates suggest different mechanism interplay between MDMs and PMNs against BCG and Map

Correlation analyses were performed between cytokine levels, and mycobacteria killing rates to evaluate the strength of relationship between both parameters (Table 1). The correlation study was performed including the data from all the culture types together and then subdivided by culture type and stimuli. IL-1β and IL-8 levels were positively correlated at 24h including all culture types in the analysis indicating that both cytokines are secreted in the same culture conditions in our assay. This correlation improved when eliminating the CCs from the analysis suggesting that cytokine release follows a different pattern in CCs maybe due to the difference in cell number. BCG-stimulated cultures showed a positive correlation between IL-1β and killing rate, which increased when eliminating CCs from the analysis. On the other hand, Map-stimulated cultures showed a negative correlation between IL-1β and killing rates, and in this case, power was increased when analysing PMNs alone. IL-8 and killing capacity were positively correlated in MDM cultures. IL-8 and killing rates were negatively correlated in Map-stimulated cultures.

**Table 1.** Correlation analyses results between cytokine levels and mycobacteria killing rates

| Culture |  |  | Stimuli |  | Parameters |  |  | n | Correlation |  |
| --- | --- | --- | --- | --- | --- | --- | --- | --- | --- | --- |
| PMN | MDM | CC | BCG | Map | IL-1β | IL-8 | killing | | coefficient | p |
| ✓ | ✓ | ✓ | ✓ | ✓ | ✓ | ✓ |  | 24 | 0.64 | <0.0001 |
| ✓ | ✓ |  | ✓ | ✓ | ✓ | ✓ |  | 16 | 0.75 | 0.0011 |
| ✓ | ✓ | ✓ | ✓ |  | ✓ |  | ✓ | 12 | 0.75 | 0.0074 |
| ✓ | ✓ | ✓ |  | ✓ | ✓ |  | ✓ | 8 | 0.95 | 0.0011 |
| ✓ | ✓ |  |  | ✓ | ✓ |  | ✓ | 12 | -0.7 | 0.011 |
| ✓ |  |  |  | ✓ | ✓ |  | ✓ | 4 | -0.98 | 0.016 |
|  | ✓ |  | ✓ | ✓ |  | ✓ | ✓ | 8 | 0.87 | 0.0044 |
| ✓ | ✓ | ✓ | ✓ | ✓ |  | ✓ | ✓ | 12 | -0.8 | 0.002 |

## 4 Discussion

In the present study, we compared aspects of the interaction of bovine PMN and MDM *in vitro* with Map and *M. bovis* BCG, a better studied mycobacterium. Specifically, we assessed whether a cooperation of both innate immune cell types would be advantageous for the host in eliminating these mycobacteria by analysing ET release, pro-inflammatory cytokine levels and killing of mycobacteria. To the best of our knowledge, this is the first study describing ET release against Map.

Bovine PMNs and MDMs were cultured separately or together. PMNs showed higher ET release against killed mycobacteria than against live bacteria as well as in comparison to MDM (Figure 2). This is in line with other observation that live mycobacteria contains glycolipids that act as antagonist to TLR2, thus inhibiting activation of innate immune cells, whereas heat-inactivation allowed for recognition by TLR2 and Mincle (35). Interestingly, activation of Mincle expressed on PMN leads to NET formation (36). In contrast, direct interaction of MDM and PMN resulted in less ET release compared to PMN cultured on their own or PMN cultured in a transwell system (see Figure 3). Indeed, this was observed for all direct co-cultures tested, independent of the stimuli tested and independent whether these were alive or inactivated. This generalized lower ET detection in direct co-cultures could be due to ET removal by efferocytosis, a process in which apoptotic cells (PMNs and/or MDMs) are removed by phagocytic cells (MDMs), to the dampening of PMN response by MDM, or to the activation of PMN by MDM resulting in other effector mechanisms than the release of ET.

Thus, we believe that in situations of direct cell-to-cell contact, ET release would not be the principal mechanism exerted by PMN, and this would be in line with our observation that PMN in co-culture situation (CC) present more internalized BCG bacilli and less detectable ETs. In contrast, in Map stimulated CCs, PMNs appear to have condensed nuclei indicative of apoptosis and no detectable histones, rarely detectable ETs and few internalized Map bacilli, whereas MDMs have histones in their cytoplasm. Whether this is a direct effect of Map on PMN, or a result of the co-culture to enable macrophages to take up and present PMN remains to be investigated. Indeed, histones can be actively secreted into the extracellular space by activated inflammatory cells (37) and by ET release (38) or are derived more passively by apoptotic and necrotic cells (37). Inhibition of efferocytosis by extracellular histones has been reported in both *in vitro* and *in vivo* conditions (39). In our experimental conditions, we believe that in Map stimulated CCs, activated or apoptotic PMNs liberate histones alone or with ETs that are engulfed subsequently by MDMs, inhibiting efferocytosis and for this reason apoptotic PMNs are still present but ETs are not detectable.

With regards to the differences seen for bacterial killing, these were most pronounced against both mycobacteria in MDM alone. MDM were highly efficient in killing the attenuated *M. bovis* BCG, but they seemed to be unable to kill Map under the tested conditions. In contrast, PMNs showed a similar killing capacity against both mycobacteria species tested. Interestingly though, when both immune cell types were co-cultured in a 1:1 ratio, the killing capacity was unaltered with regards to BCG, but was similar to that obtained in PMNs for Map. Nevertheless, taking into account that CCs are composed by half of each cell type compared to MDM or PMN cultures, the obtained CC results indicate that a synergistic effect (observed killing > expected killing) in terms of improved killing capacity in BCG-infected co-cultures is observed, whereas in Map-infected co-cultures the effect was additive (observed killing = expected killing). Thus, the presence of PMN at the site of infection during the initial stages of Map may be beneficial for the resolution of the disease. Indeed, several studies (40)(41) demonstrated that human PMN are capable of controlling Mtb infection *in vitro* and murine PMN have been shown to kill *Mycobacterium avium ex vivo* when activated *in vivo* with G-CSF (42).

In contrast to our data regarding ET formation and killing, MDM produced higher cytokine levels compared to PMNs and CCs in response to both mycobacteria. Cytokine levels of co-cultures were lower than expected, resulting equivalent to those obtained in cultures of PMNs alone. A study by Sawant and McMurray showed an additive effect of IL-8 and IL-1β release in co-cultures of guinea pig PMN with alveolar MΦ challenged *in vitro* with Mtb (43). It is likely that those results were obtained because their co-cultures harboured PMN and MΦ in a 1:1 ratio, but doubling the total amount of cells, whereas our study halves the amount of each cell type to preserve the total cell number. In addition, differences on host and mycobacteria species and MDM sources (lung isolated cells in their case and peripheral isolated cells in our case), make a comparison between both studies difficult.

However, it is also possible that anti-inflammatory mediators released by PMN act in an autocrine/paracrine manner, suppressing the production of pro-inflammatory cytokines by MDM, similar as described before (44). Indeed, the high IL-1β and IL-1α levels described by others in PTB (45) could be a result of the activation and recruitment impairment of PMN (28, 29; 47–49) and the resulting low numbers of PMN associated to PTB granulomatous lesions (50, 51). Moreover, high levels of these cytokines are proposed as candidates responsible for PTB inflammation process and contributors to the development of the Th17 response during the final stages of the disease (50).

Similarly, IL-8 expression has also been described to increase in PTB and monocytes stimulated with Map (53, 54) compared to controls. In our study IL-8 secretion of the same individuals against BCG and Map was compared. The lower IL-8 release of Map versus BCG stimulated MDMs could be a reflection of higher pathogenicity of Map in MDMs. In line with this observation is the fact that multi-drug resistant Mtb strains decreased release of IL-8 by infected bronchial epithelial cells, thus limiting PMN recruitment to the site of infection (53).

When correlating cytokine release with killing, we observed a positive correlation with the attenuated *M. bovis* BCG strain, being higher in MDM, whereas the response to Map infection was negatively correlated with killing, showing a stronger effect in PMN. Low IL-8 levels have been associated to poor prognosis in hTB (23) and survival of *Mtb* inside MΦ by altering PMN effector functions (53). In the present experiments, IL-8 levels were high in PMN, MDM and co-cultures, indicating that despite the negative correlation with Map killing, more factors are highly likely to be involved in this interaction. As for IL-1β, previous studies in IL-1 knock-out mice showed an increased susceptibility to Mtb infection, developing a higher bacterial burden and mortality (54). With this in mind, we would have expected that the lowest mycobacterial survival rate would be seen in the cultures with the highest levels of IL-1β. Whereas this was the case for BCG, it was not the case of Map-stimulated MDM cultures, indicating once again that more factors are involved. In fact, *in vitro* studies indicate that Map infection promotes a self-destruction state in the epithelium caused by increased IL-1β levels that attract MΦ, thus providing Map with an escape route from destruction (55).

Based on the results obtained in our experiments, we hypothesize that bovine PMN may have an important role against Map. PMN trigger inflammatory responses through IL-1β release and start an effective response phagocytosing and immobilizing bacteria with their NETs. As a consequence, MΦ are activated and attracted by IL-1β production to the site of infection. Upon arrival, MΦ would phagocytose neutrophils’ ETs and apoptotic rests, which provides them with the antimicrobial compounds produced by neutrophils after first contact with Map. This efferocytosis activity, as part of the homeostasis process, would reduce the inflammatory response leaded by IL-1β, IL-23 and IL-17 expression, ending ultimately in Map destruction. The timing is probably important in this cooperation and if MΦs arrive at the infection site too early, PMN might not have enough time to effectively expand their antimicrobial mechanisms, MΦ will be the dominant cell type attempting Map elimination but end-up being taken over by Map as a Trojan horse. PMN have been postulated as innate effectors of TB resistance due to their highly effective antimicrobial effector mechanisms they can expand during the early stage of Mtb infection (56). Applied to Map, it could be that harbouring a particular number of competent and tightly regulated PMN capable of expanding the correct antimicrobial mechanism at the specific time may be the difference between being resistant to PTB or developing subclinical or clinical disease.

## Conclusions

Taking all assayed parameters into consideration, we could conclude that cooperation of MDMs and PMNs may confer advantages through different mechanisms depending on the pathogen. As for BCG, co-cultures show good levels of elimination of the mycobacteria (high killing rates) and lower cytokine levels and ET release than the PMN and MDM cultures, which, could translate to a decreased tissue damage in the living organism, since an excessive inflammatory reaction would not take place. In Map stimulated cultures, co-cultures show slightly worse figures for mycobacteria killing compared to PMNs but improved rates compared to MDMs, also with lower levels of pro-inflammatory cytokines. In BCG, ET release is negatively correlated to bacterial killing, suggesting that killing could be due to phagocytosis, whereas as in Map ET release is positively correlated to bacterial killing, suggesting that probably most killing is due to PMNs ETosis and provision of macrophages with killing capacity molecules in the co-cultures. Further research should involve *in vivo* studies to confirm that PMNs are necessary in the defence against Map and studies clarifying which subsets of PMNs are involved in each effector mechanism in order to develop therapeutic agents targeted at improving PMN competence.

## Author Contributions

DW and NE conceived the study. AU carried out the animal sampling. IL, EM, AH, JK, HH and NE performed the laboratory work. IL and NE compiled and analyzed the data. IL and NE collated the results. IL, NE, JA and DW drafted the preliminary manuscript. All authors participated in the review and the editing of the final draft and also read and approved its final version.

## Acknowledgments

The authors want to thank Ainara Badiola and Maddi Oyanguren for technical support and Joseba Garrido for critical reading of the manuscript. Funding was provided by Spanish central government and Basque research project PROBAK (RTA 2017-00089-00-00) and by the Departamento de Economía e Infraestructuras of the Basque Government. IL held a pre-doctoral grant from Departamento de Economía e Infraestructuras of the Basque Government (2017) and was granted an EMBO short-term fellowship (8407) and a FEMS research and training grant (FEMS-GO-2019-507). The funders had no role in the study design, data collection and interpretation, or the decision to submit the work for publication.

## Data Availability Statement

The datasets generated for this study are available on request. The raw data supporting the conclusions of this article will be made available by the authors, without undue reservation.

